# IbpAB small heat shock proteins are not host factors for bacteriophage ϕX174 replication

**DOI:** 10.1101/2022.10.13.511849

**Authors:** Hannah X Zhu, Bradley W Wright, Dominic Y Logel, Mark P Molloy, Paul R Jaschke

## Abstract

Bacteriophages exploit host proteins for successful infection. Small heat shock proteins are a universally conserved family of stress-induced molecular chaperones that prevent irreversible aggregation of proteins. Two small heat shock proteins, IbpA and IbpB, are a class of holding modulators or “holdases”, which bind partially folded proteins and await ATP-driven folding chaperones for refolding. Bacteriophage ϕX174 is a small, icosahedral, and non-tailed virus belonging to the *Microviridae*. During ϕX174 infection of *Escherichia coli* C122, IbpA and IbpB were previously found to be the most highly upregulated host proteins, with expression levels comparable to ϕX174 proteins. In this work, to understand the role of IbpA and IbpB during ϕX174 infection, we used a hybrid approach of CRISPR interference and genomic knockouts to disrupt the *ibpA* and *ibpB* genes. We show that these two proteins do not appear to be necessary for efficient ϕX174 replication, and moreover, their absence has no effect on ϕX174 fecundity.

**Importance:** The small heat shock proteins (sHsps) are universally conserved family of stress-induced molecular chaperones that prevent irreversible protein aggregation. In *E. coli*, the IbpA/B sHsps work together, and separately, to bind partially folded proteins and await ATP-driven folding chaperones for refolding. These proteins are highly upregulated during protein overexpression and bacteriophage infection, but their collective role in bacteriophage infection has not been investigated. Here, we show that the *ibpA/B* genes are dispensable for bacteriophage ϕX174 infection, and are likely not essential host factors despite their abundance during diverse phage infections. Instead, this work points towards their role as cell wall integrity sensors, similar to the phage shock protein system, in addition to their canonical role as holdases of cytoplasmic protein.

## Introduction

Bacteriophages (phages) are viruses that obligately infect and replicate within a bacterial host. Phages are the most abundant entities within the biosphere, with an estimated global population of ~ 10^31^ individual phage particles that outnumber their hosts by orders of magnitude (1–3). Phage-host interactions represent a vital ecological component of microbial systems with an estimated 10^23^ viral infections occurring per second in the global oceans (4). Genomic knockout studies and genome-wide screens have revealed that successful phage infections rely on diverse bacterial host factors, including phage receptors and regulators, lipopolysaccharide (LPS) biosynthesis genes, and DNA replication and transcription machinery components (5–9). Among all identified host factors, the heat shock proteins (Hsp) are one of the most upregulated host factors during phage infections, at both transcription and translation levels (10–12).

Hsp are a class of molecular chaperones and are produced by bacteria under challenging conditions such as temperature, oxidative, or osmotic stresses. Their fundamental role is to facilitate the folding of unfolded and partially folded proteins through ATP-driven binding and releasing proteins in their functional states (13, 14). Several Hsps have been deemed essential to the development of viable phage virions. Phage λ requires the *oriλ*-O-P-DnaB nucleoprotein structure to form an initiation complex for DNA replication; the Hsps DnaK, DnaJ, and GrpE (DnaKJE) release P protein from the λ initiation complex, and ClpX degrades O protein, freeing DnaB helicase to unwind the phage DNA (15–17). The Hsp DnaJ, independently facilitates the excision of prophage KplE1 of *Escherichia coli* K12 through interactions with the recombination directionality factor TorI (18). DnaJ has also been found to enhance the lysis of host cells by *E. coli* phage MS2 via the formation of a membrane-associated complex with lysis protein L (19). Additionally, the GroEL/GroES chaperonins assist with phage capsid morphogenesis by facilitating the correct folding of coat proteins (20–22).

Bacteriophage ϕX174 is a small, icosahedral, and non-tailed virus belonging to *Microviridae* (23) and work has shown that mutations in DnaJ/DnaK/GrpE/GroEL/GroES make the *E. coli* host more sensitive to lysis by E protein (24). In addition to this work, we have recently shown that the small heat shock proteins (sHsps) IbpA and IbpB were the most upregulated host proteins during ϕX174 infection, with levels matching or exceeding phage protein, which suggests they are important for efficient phage infection (10).

The sHsps are a widespread and diverse family of low-molecular weight Hsps with monomeric mass of 12–43 kDa. Structurally, sHsps share a highly conserved α-crystallin domain (ACD) (~ 90–100 amino acid residues) that is required for the formation of sHsp homodimer, which is the building block of higher oligomers, flanked by variable and disordered N- and C-terminal extensions (25, 26). sHsps have been extensively studied for their role as “holdases” to prevent irreversible and uncontrolled aggregation of proteins, in an ATP-independent manner. Some sHsps also exhibit sequestrase activity that facilitates refolding of proteins in near-native state, and cytoprotective aggregase activity through the formation of deposits of misfolded proteins (25). sHsps are conserved across all domains of life. The dissemination of sHsps across all domains of life and the fact that genomes often contain multiple paralogs is indicative of their importance to cells, and has been discussed in detail (25, 27).

In *E. coli*, IbpA and IbpB are organised under a single operon (*ibpAB*) within the σ^32^ regulon and the independent regulation of *ibpB* within the σ^54^ regulon. These proteins have been found specifically associated with inclusion bodies and intracellular aggregates of overexpressed proteins during both heat shock and recombinant protein expression (28–30). Following heat-induced transcription of *ibpAB* operon, the bicistronic transcript is processed into monocistronic *ibpA* and *ibpB* transcript; the translation of *ibpA* and *ibpB* is controlled by an RNA thermometer in its 5’ untranslated region, forming a secondary structure that blocks the binding of 30S ribosomal subunits at low temperature (31). It has also been found that oligomeric IbpA suppresses translation of its own mRNA transcript as well as that of *ibpB* (32). Collectively, *ibpA* and *ibpB* genes are essential when cells experience prolonged heat stress, > 50°C for 4 hours (33). IbpA and IbpB form oligomeric assemblies with partially folded proteins and deliver the substrates to the ATP-driven DnaKJE-ClpB bi-chaperone network, for refolding (34–38). IbpA and IbpB are known to act cooperatively, with IbpA stably interacting with aggregating substrates, preventing inter- and intra-molecular hydrophobic interactions by reducing the size of IbpAB-substrate assembly, while IbpB is required for the dissociation of the bound substrate from the assembly for further processing (35, 36, 39, 40). Despite this cooperation and differentiation of function, the loss of either chaperone alone results only in a minimal disruption to cell growth (41), as they are thought to be functionally redundant (42), while DnaKJE-dependent refolding of denatured proteins would still be expected to occur in absence of either IbpA or IbpB, although at a considerably lower rate and efficacy (43, 44). Because of this known functional redundancy and genome-wide screens showing *individually* disrupted *ibpA* and *ibpB* genes were non-essential for phage replication (e.g., T4, T7, and λ phages of *E. coli*) as well as not conferring phage resistance (5–7, 9, 45), we suspected their role in phage replication could be functionally redundant and warranted further investigation.

In our previous work, the IbpA and IbpB were the most highly upregulated of any host protein during bacteriophage ϕX174 infection of *E. coli* C122 (10). Thus, we reasoned that these host proteins could either be a defense mechanism deployed by the cell to attempt to sequester the phage proteins in an unfolded state and prevent capsid formation, or that these proteins were an important host factor that enabled large numbers of ϕX174 capsid proteins to be held in a nearly folded state to prevent the formation of inclusion bodies, and then assembled efficiently into the capsid when needed in the replication cycle.

In this work we determined the role of IbpA/B in the ϕX174 infection cycle using a combination of CRISPR gene knockouts and CRISPRi knockdowns, and demonstrate that they are collectively non-essential for ϕX174 replication. Our results suggest that IbpA and IbpB are neither a host phage defense factor nor an essential host factor required for viral replication.

## Results

### IbpA/B are functionally redundant for ϕX174 replication but not for heat stress tolerance

To determine the effect of these proteins on bacteriophage ϕX174 replication we created single knockouts of *ibpA* and *ibpB* in the *E. coli* C122 strain using CRISPR-Cas9 (Table 1). We designed the gene knockouts which resulted in highly truncated IbpA (11% of wild-type) and IbpB (31%) proteins by incorporating pre-mature stop codons into the coding sequence (Fig. 1A). We confirmed these knockouts through whole genome sequencing. As expected from previous work (29, 32, 42, 46), both C122Δ*ibpA* and C122Δ*ibpB* strains displayed no growth defects in rich liquid media at 37°C. However, growth defects at 45°C were apparent (Fig. 1B, C and D).

**Table 1.** Description of Plasmids, Strains, and Phage Used in this Study

| Name | Description | Reference or source |
| --- | --- | --- |
| **Plasmids** |  |  |
| pULTRA-CNF | Plasmid used for pULTRA complement construction. Contains spectinomycin resistance marker | Addgene: # 48215 (A gift from Peter Schultz Lab) (48) |
| pULTRA::*ibpAB* | Plasmid containing native C122 *ibpAB* operon (coordinates 16,177-17,263 from NZ_LT906474). Contains spectinomycin resistance marker | This study |
| pULTRA::empty | Plasmid used as pULTRA control, without insert, containing spectinomycin resistance marker | (49) |
| pKDsgRNA-ack | Plasmid used for cloning guide RNA (gRNA). Contains spectinomycin resistance marker, temperature sensitive origin of replication (cure at 37°C), and lambda red recombinase machinery under arabinose inducible promoter | Addgene: # 62654 (A gift from Kristala Prather Lab) |
| pKDsgRNA-16400 | gRNA plasmid targeting *ibpA* gene. Contains spectinomycin resistance marker, temperature sensitive origin of replication (cure at 37°C), and lambda red recombinase machinery under arabinose inducible promoter | This study |
| pKDsgRNA-16890 | gRNA plasmid targeting *ibpB* gene. Contains spectinomycin resistance marker, temperature sensitive origin of replication (cure at 37°C), and lambda red recombinase machinery under arabinose inducible promoter | This study |
| pKDsgRNA-p15 | gRNA plasmid targeting pCas9-CR4 plasmid. Enables curing. Contains spectinomycin resistance marker, temperature sensitive origin of replication (cure at 37°C), and lambda red recombinase machinery under arabinose inducible promoter | Addgene: # 62656 (A gift from Kristala Prather Lab) |
| pCas9-CR4 | Contains Cas9 nuclease gene under control of pTet promoter with *ssrA* tag and constitutive *tetR* | Addgene: # 62655 (A gift from Kristala Prather Lab) |
| pFR56::ns | Plasmid used for cloning gRNA. Contains chloramphenicol resistance marker, a control non-specific single gRNA (sgRNA) under a constitutive promoter, and *dcas9* under the control of a 2,4-diacetylphloroglucinol (DAPG)-inducible PhIF promoter | GenBank accession no. MT412099 (A gift from David Bikard Lab) |
| pFR56::*ibpA* | gRNA plasmid targeting *ibpA* gene. Contains chloramphenicol resistance marker, sgRNA under a constitutive promoter, and *dcas9* under the control of a DAPG-inducible PhIF promoter | This study |
| pFR56::*ibpB* | gRNA plasmid targeting *ibpB* gene. Contains chloramphenicol resistance marker, sgRNA under a constitutive promoter, and *dcas9* under the control of a DAPG-inducible PhIF promoter | This study |
| **Strain** |  |  |
| *Escherichia coli* C122 | Wild-type host strain of bacteriophage ΦX174. | National Collection of Type Cultures (NCTC) strain NCTC122 |
| C122(pULTRA::empty) | C122 strain harbouring pULTRA:: empty | This study |
| C122(pULTRA::*ibpAB*) | C122 strain harbouring pULTRA::*ibpAB* | This study |
| C122Δ*ibpA* | C122 strain harbouring genomic disruptions of *ibpA* | This study |
| C122Δ*ibpB* | C122 strain harbouring genomic disruptions of *ibpB* | This study |
| C122Δ*ibpAB* | C122 strain harbouring genomic disruptions of *ibpA* and *ibpB* | This study |
| C122Δ*ibpAB* (pULTRA::empty) | C122 strain harbouring genomic disruptions of *ibpA* and *ibpB* and containing pULTRA::empty plasmid | This study |
| C122Δ*ibpAB* (pULTRA::*ibpAB*) | C122 strain harbouring genomic disruptions of *ibpA* and *ibpB* and containing pULTRA::*ibpAB* plasmid | This study |
| C122(pFR56::ns) | C122 strain harbouring pFR56::ns plasmid targeting a loci not present within *E. coli* | (7) |
| C122(pFR56::*ibpA*) | C122 strain harbouring pFR56 plasmid targeting *ibpA* | This study |
| C122(pFR56::*ibpB*) | C122 strain harbouring pFR56 plasmid targeting *ibpB* | This study |
| C122Δ*ibpA*(pFR56) | C122 strain harbouring genomic *ibpA* knockout and pFR56 plasmid | This study |
| C122Δ*ibpB*(pFR56) | C122 strain harbouring genomic *ibpB* knockout and pFR56 plasmid | This study |
| C122Δ*ibpA*(pFR56::*ibpB*) | C122 strain harbouring genomic *ibpA* knockout and pFR56 plasmid targeting *ibpB* | This study |
| C122Δ*ibpB*(pFR56::*ibpA*) | C122 strain harbouring genomic *ibpB* knockout and pFR56 plasmid targeting *ibpA* | This study |
| **Bacteriophage** |  |  |
| Bacteriophage ΦX174 | Phage of *E. coli* C synthesized to be identical to Sanger GenBank sequence NC_001422.1 | (50) |

**FIG 1.**
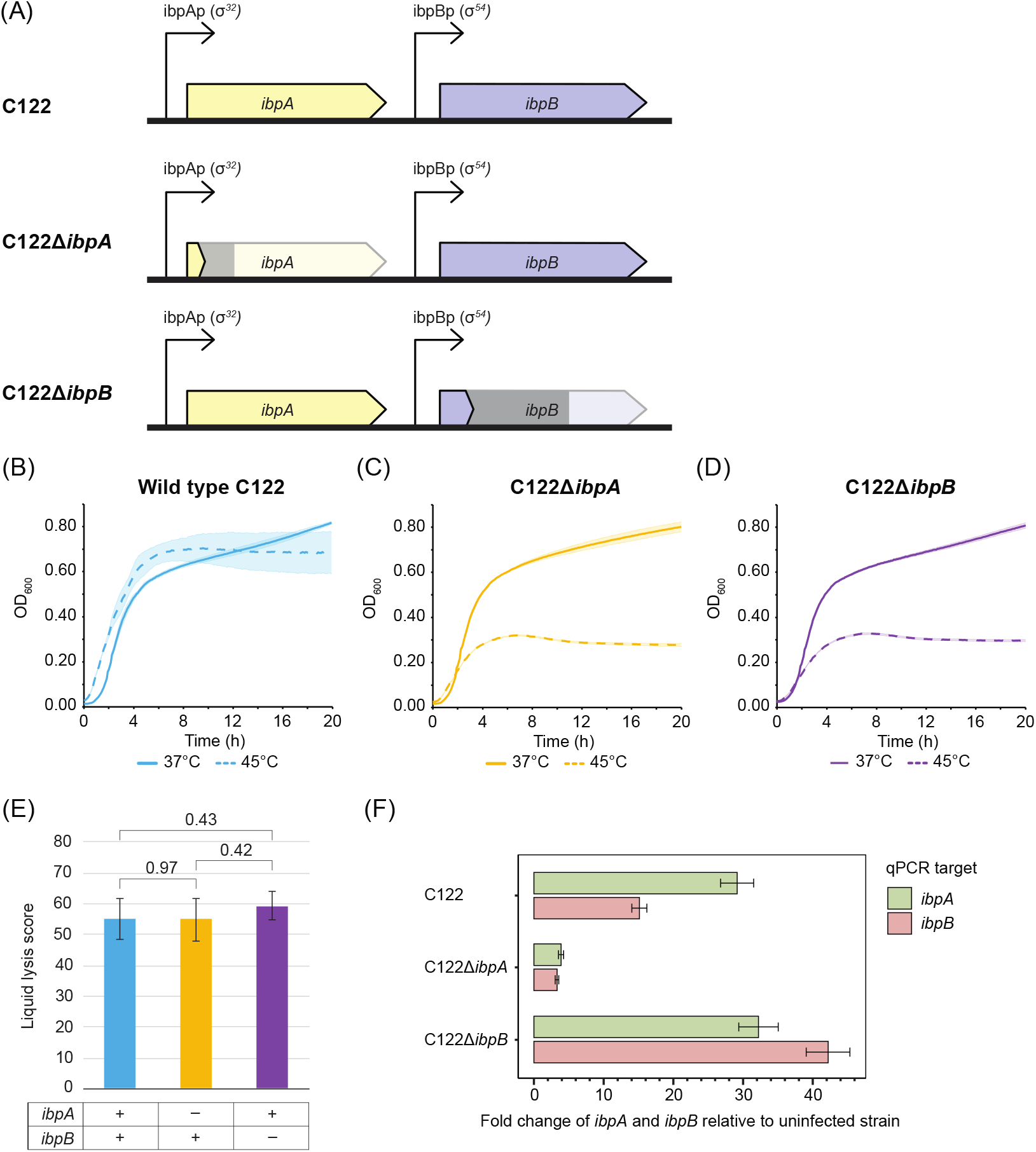
Single *ibpA* and *ibpB* gene disruptions result in growth inhibition at elevated temperatures but no difference in ϕX174 replication efficiency. (A) *ibpAB* gene knockout scheme. Wild-type *ibpA* is 355 bp long while knockout introduces premature stop codon resulting in coding sequence of only 39 bp (11%). Wild-type *ibpB* is 233 bp long while knockout introduces premature stop codon resulting in coding sequence of only 77 bp (31%). Grey bars indicate area of each gene removed after repair and faded gene symbol indicates wild-type extent of gene. (B-D) Growth rate comparison between 37°C and 45°C in *ibpA* and *ibpB* knockout strains. (E) Comparison of ϕX174 virulence against wild type C122, C122Δ*ibpA*, and C122Δ*ibpB* bacterial strains. The genetic background of the *E. coli* host denoted by + (wild-type gene present) or – (gene disrupted). Phages were added to exponential-phase cultures at an MOI of 5 and bacterial cultures were grown for a total of 24 h with OD600 measured every 5 min. Absorbance changes over time were compared to a non-infected control culture. The liquid assay score is the difference in the area under the growth curves of the phage-infected sample and the non-infected control (47). No inhibition of bacterial growth results in a score of zero, whereas complete of absence of growth gives a score of 100. Brackets with numbers above refer to Student’s two-tailed t-test p-values. (F) RT-qPCR measurement of *ibpA* and *ibpB* transcript targets comparing the expression of the target genes when infected to the uninfected strains to determine the expression induction.

To determine what effect the loss of one of the sHsps had on ϕX174 replication, we infected C122Δ*ibpA* and C122Δ*ibpB* strains along with wild-type C122 and measured changes in OD_600_ over time. The results showed that infections in the single-gene knockout strains were indistinguishable from wild-type *E. coli* at MOI 5 (Fig. 1E). A reduced MOI 0.001 infection also resulted in similar results between single-gene *ibpA* or *ibpB* knockouts and wild-type (<u>Supplemental Fig. S1</u>), but didn’t induce *ibpAB* strongly, and thus MOI 5 was used going forward. Measuring the disruption effects of the *ibpA* and *ibpB* knockout gene sequences on transcript abundance of the *ibpAB* operon using RT-qPCR showed that the *ibpA* disruption appears to impair expression of both *ibpA* and *ibpB* transcripts when cells are infected with ϕX174, while the *ibpB* disruption does not (Fig. 1F).

We reasoned that even with the low transcriptional induction of the *ibpA* and *ibpB* transcripts in C122Δ*ibpA*, there could still be functional IbpB protein produced at a level that could mask any beneficial or deleterious effects on ϕX174 replication (Fig. 1E).

We next attempted to create a double knockout of the *ibpAB* genes using CRISPR/Cas9 in a similar manner to the single-gene disruptions. We found our initial attempts to create this strain resulted in obtaining transformants with off-target effects that fell into two categories: (1) cryptic recombinants restoring the *ibpB* gene in screened colonies (Fig. 2A), or (2) apparently successful disruption of both genes but with off-target mutation elsewhere in the genome.

**FIG 2.**
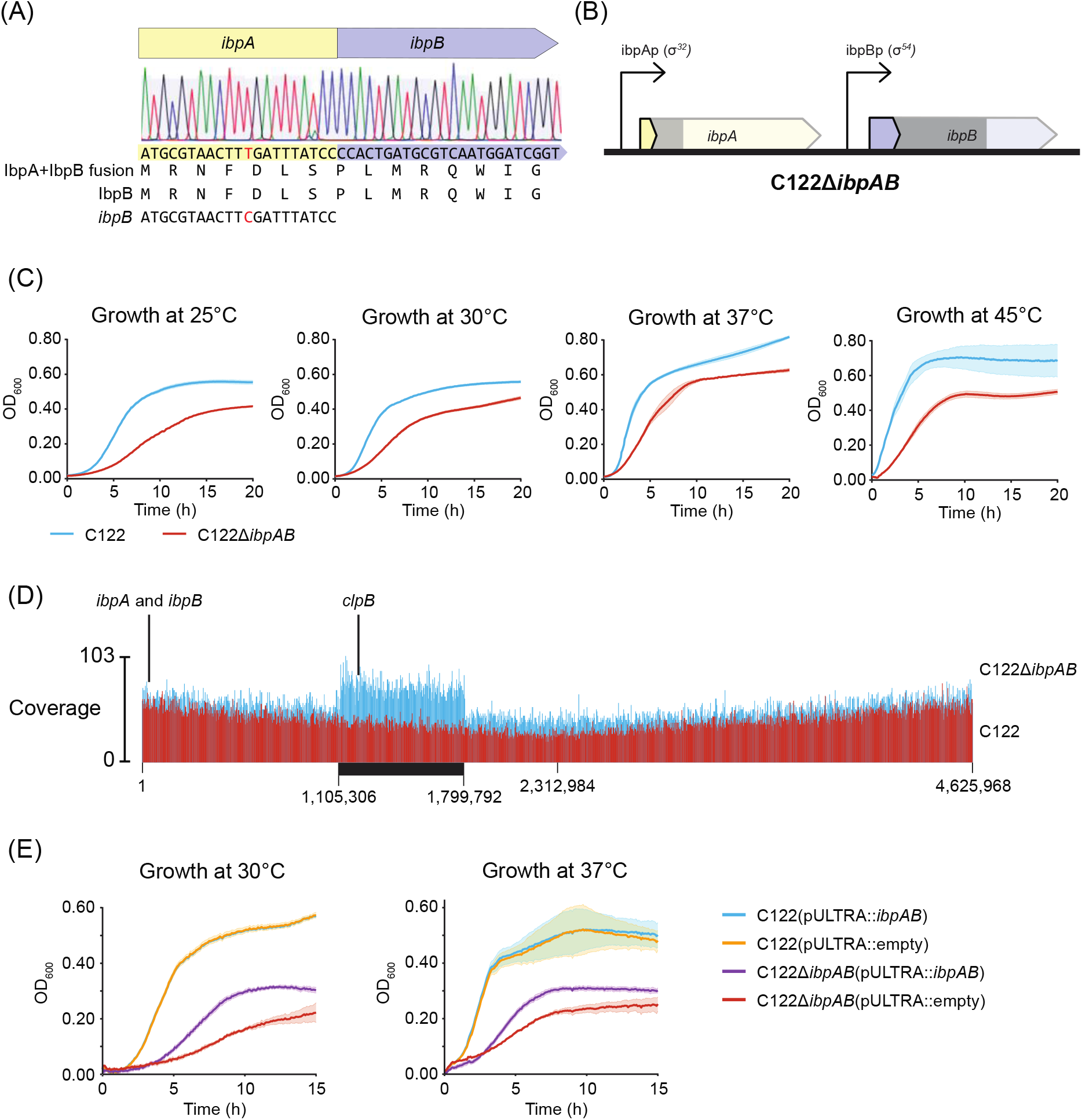
Difficulty obtaining a double *ibpAB* knockout without off-target effects. (A) Cryptic recombination between *ibpA* 5’-end and *ibpB* gene 3’-remnants creates *ibpB* gene with identical amino acid sequence to IbpB protein. (B) Apparent double ibpAB disruption where a premature stop codons were introduced into *ibpA* and *ibpB* genes resulting in coding sequences of only 39 bp (11%) and 77 bp (31%) of the wild-type genes, respectively. Grey bars indicate area of each gene removed after repair and faded gene symbol indicates wild-type extent of gene. (C) Apparent double-knockout *ibpAB* strain growth characteristics at 25° - 45°C compared to isogenic wild-type C122 strain. (D) Deep sequencing coverage of C122*ΔIbpAB* strain (blue) compared to wild-type C122 (red) reveals putative 694.5 kb sequence duplication outside *ibpAB* region cut by CRISPR/Cas9. Black bar indicates region of increased read coverage with approximate genomic coordinates of boundaries. (E) Complementation of C122*ΔIbpAB* strain with plasmid-borne *ibpAB* genes shows only partial restoration of growth defect at 30°C and 37°C. Plasmid pULTRA::*ibpAB* denotes pULTRA harboring native *ibpA/B* operon while pULTRA::empty denotes empty pULTRA plasmid backbone. Variance of one standard deviation (n = 3) in measurements shown as faded region of same colour around growth curves.

In a strain where we confirmed that both *ibpA* and *ibpB* genes were disrupted (Fig. 2B), we found that it displayed a severe a growth defect at 25°C, 30°C, 37°C, and 45°C (Fig. 2C). Whole genome sequencing of this strain revealed that an area of ~694.5 kb long sequence resulted in approximately twice the read coverage of adjacent areas of the C122Δ*ibpAB* sequence and of the corresponding area of the C122 genome sequence (Fig. 2D). Furthermore, this putatively duplicated region was present outside of the *ibpAB* gene region target by the CRISPR guide RNAs (Fig. 2D) and contained 606 coding sequences, seven ncRNAs, tmRNA, three rRNAs, and 10 tRNAs, thus making the precise assessment of any epistatic effects unlikely. One coding sequence known to be involved in the same pathway as *ibpAB* that was present within this genomic region was the *clpB* gene, the product which in cooperation with DnaK uses energy derived from ATP hydrolysis to dissociate the denatured protein aggregates from sHsp-substrate complex (51). Complementation of the C122Δ*ibpAB* strain with plasmid-borne *ibpAB* genes failed to fully restore the growth phenotype (Fig. 2E) and thus we did not investigate this strain further.

Due to the difficulty of obtaining a clean double *ibpA* and *ibpB* knockout on the C122 strain genome, and all previously made double *ibpAB* knockouts were in *E. coli* strain backgrounds that are not susceptible to ϕX174 infection (52), we chose to deploy CRISPR interference (CRISPRi) in the single C122Δ*ibpA* and C122Δ*ibpB* knockout backgrounds to create the same effect with an inducible knockdown. CRISPRi employs a catalytically dead Cas9 (dCas9), which forms a complex with a single-guide RNA (sgRNA) (consisting of a gRNA and a structural scaffold) and is directed to the target gene to silence its expression (53, 54). We designed gRNAs that targeted the coding strand of *ibpA* and *ibpB* genes because they more efficiently repress transcription than noncoding strand targets (53, 54). The target region for the sgRNAs was within the first 100 nucleotides of the CDS and was upstream of the sequence predicted to be the protein active site (55), such that translation of the incomplete mRNA transcript (if any) would result in a non-functional truncated protein.

The CRISPRi expression plasmid, pFR56, contains a constitutively expressed sgRNA and a *dcas9* gene controlled by a 2,4-diacetylphloroglucinol (DAPG)-inducible PhlF promoter (Fig. 3A) (7). We assembled pFR56 with gRNA sequences targeting either *ibpA* or *ibpB*, and created CRISPRi expression vectors pFR56::*ibpA* and pFR56::*ibpB* that suppress the transcription of *ibpA* and *ibpB*, respectively.

**FIG 3.**
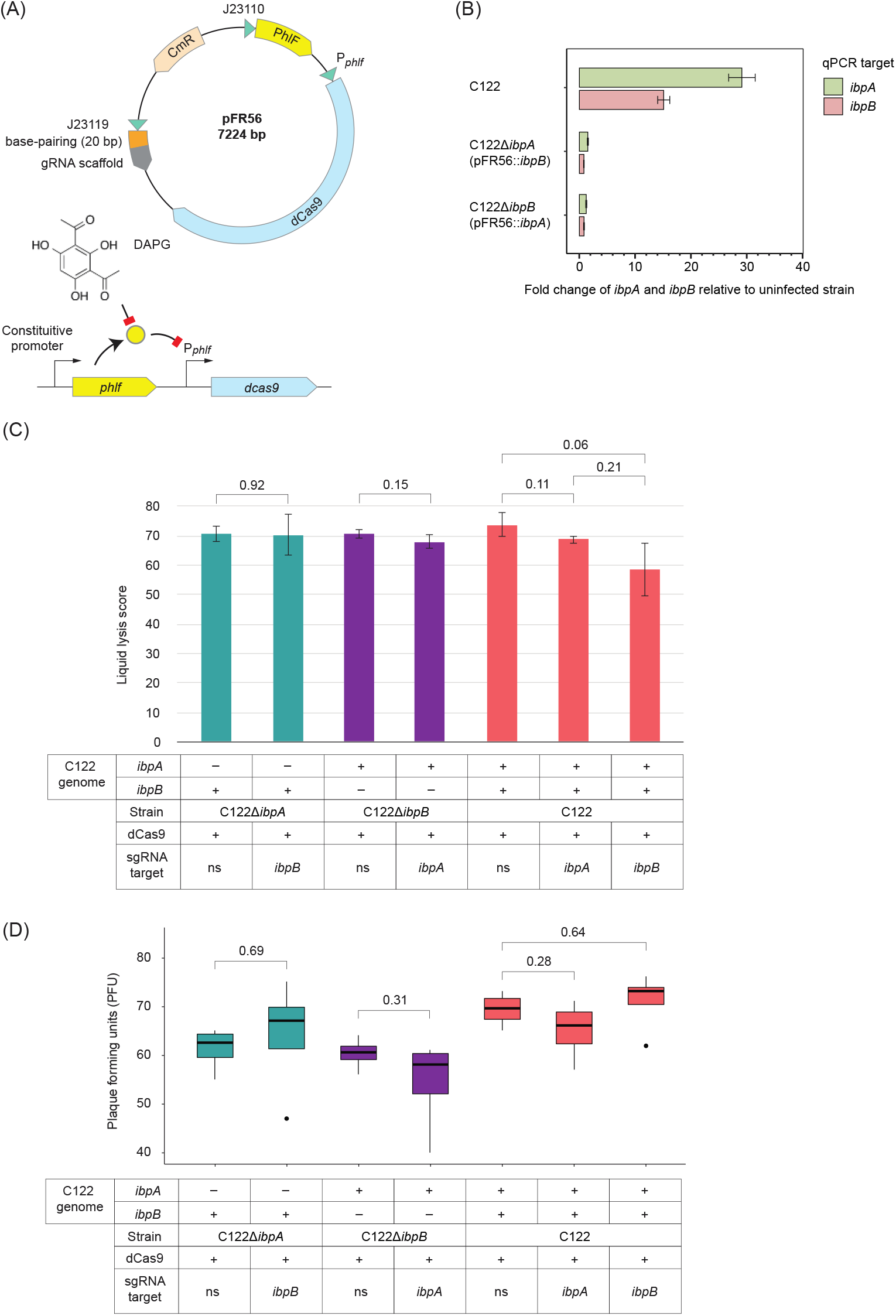
CRISPRi-mediated *ibpAB* disruption shows no effect on ϕX174 infection. (A) CRISPRi system. pFR56 with constitutively expressed sgRNA and the *dcas9* gene under control of DAPG-inducible PhlF promoter (7). (B) RT-qPCR analysis of CRISPRi-mediated knockdown strains infected with ϕX174. CRISPRi-mediated *ibpAB* knockdown strains were grown in phage-LB at 37°C/240 rpm to early exponential phase when dCas9 was induced by DAPG (50 µM). At 40 min after induction, the bacteria were infected with wild type ϕX174 at an MOI of 5. Cells were harvested at 60 min post-infection, and RNA was extracted. RT-qPCR against *ibpA* and *ibpB* targets was performed in biological and technical triplicate. (C) ϕX174 liquid culture virulence against CRISPRi-mediated *ibpAB* knockdown strains. The *ibpAB* sgRNAs are named for their target and ns (non-specific) being an sgRNA that lacks a specific target. (D) ϕX174 efficiency of centre of infection against CRISPRi-mediated knockdown. Student’s two-tailed t-test p values reported.

Using the C122Δ*ibpA* and C122Δ*ibpB* strains as background, we introduced the pFR56::*ibpB* and pFR56::*ibpA* plasmids into these strains to create C122Δ*ibpA*(pFR56::*ibpB*) and C122Δ*ibpB*(pFR56::*ibpA*) strains where CRISPRi-mediated transcriptional inhibition could be induced within the non-disrupted *ibpA* or *ibpB* coding sequence, thus effectively silencing both IbpA and IbpB protein production from both genes. We also introduced into both C122Δ*ibpA* and C122Δ*ibpB* strains a plasmid (pFR56::ns) expressing a control gRNA which does not have a specific target in the *E. coli* genome and is a control for the effect of expression of an sgRNA in the presence of dCas9 itself. Based on previous work, the non-specific sgRNA is not expected to have a phenotypic effect (7). Lastly, we transformed pFR56::*ibpA*, pFR56::*ibpB*, and pFR56::ns into wild-type C122 strain as a control.

We first confirmed the efficiency of the CRISPRi by RT-qPCR of ϕX174 infected cells. The expression of dCas9 was induced by DAPG (50 µM) at early exponential phase and we allowed 40 min for dCas9 to be produced before phage infection. We did not expect *ibpA* or *ibpB* transcripts to be expressed prior to ϕX174 infection because they are within a stress response regulon (10, 12, 56, 57) and we did not subject the cells to temperature-shock. At 60 min post-infection, we found upregulation of both *ibpA* and *ibpB* transcripts in the absence of dCas9 expression, whereas the amount of *ibpA* transcripts were less than 2-fold above non-infected samples, and *ibpB* transcripts were unchanged when dCas9 was expressed (Fig. 3B). Together, these observations suggest that the gRNAs targeting the *ibpA* and *ibpB* transcripts were able to direct dCas9 binding and prevent the majority of transcription from these loci. In particular, the C122Δ*ibpA*(pFR56::*ibpB*) strain should represent a true knockout of functional IbpA and IbpB proteins, since the *ibpB* transcript is fully blocked by CRISPRi, and the slightly induced *ibpA* transcript contains a disrupted coding sequencing, presumably resulting in a non-functional IbpA protein.

We next measured ϕX174 lysis efficiency in the absence of the IbpAB proteins by comparing liquid lysis scores of each strain undergoing *ibpA* or *ibpB* CRISPRi with the same strain expressing a nonspecific sgRNA. We found no difference between ϕX174 replication efficiencies in strains with wild-type *ibpAB* expression and those with a combination of genomic disruption and CRISPRi knockdown of the *ibpAB* genes (Fig. 3C). Using a lower MOI of 0.001 also showed no significant effect of *ibpAB* disruption on ϕX174 replication (<u>Supplemental Fig. S2</u>).

Because some infection deficiencies can be obscured in bulk culture measurements (58), we next wanted to measure the efficiency of each infection event through the efficiency of centre of infection assay. We first grew *E. coli* strains, induced CRISPRi, and infected cells as before, except in this case we washed cells following infection to remove unadsorbed phage, followed by co-plating with wild-type C122 as an indicator strain. Thus, any plaques observed from this method show the result of an initial successful infection event, but not the cumulative effect of multiple rounds of infection within cells undergoing CRISPRi. Similar to the liquid lysis assay (Fig. 3C), we did not observe significant differences in ϕX174 infection efficiency when infecting *ibpAB* knockdown strains (Fig. 3D).

## Discussion

In this work, we examined the role of two sHsps, IbpA and IbpB, during ϕX174 infection of *E. coli* C122, using a combination of single-gene genomic knockout and CRISPRi-mediated transcription inhibition. We found that the virulence and plaque-forming ability of ϕX174 was unaffected by the loss of the *ibpA* and *ibpB* genes, and thus they are not host factors for this virus. This result is unexpected given the two sHsps have been found amongst the most enriched host proteins during phage infections (10, 12, 56, 57), and leaves us with the question of why they are highly expressed during ϕX174 infection?

The IbpA and IbpB proteins seem to be important for stabilising denatured protein aggregates. They have been found massively upregulated when there is intracellular protein aggregation as a result of stresses, including elevated temperature, recombinant protein overexpression, and oxidative damage (32, 33, 59–61). The sHsps are widespread and diverse family of molecular chaperones that are found across all domains of life. Although they are not in the bacterial minimal gene set (62) nor in the *E. coli* essential gene set (7), mutations in genes encoding sHsps have been linked to several human diseases, including Charcot-Marie-tooth disease, cataracts, cardiomyopathy, and motor neuropathy (63–65).

Individual knockout studies have shown the *ibpA* and *ibpB* genes to be non-essential in *E. coli* K-12 strains. Independent single deletions of *ibpA* or *ibpB* using TraDIS (41) and CRISPR interference pools (6, 66) also show these genes are individually nonessential. Double *ibpA* and *ibpB* knockout mutants in K-12 strains have been obtained and show significant viability loss only under heat stress (> 50°C) for extended periods (> 2 h) (29, 33, 34, 39, 42). In double *ibpA* and *ibpB* knockout strains, an additional knockout of Hsp *clpB* showed increased requirements for the DnaK chaperone system for reversal of protein aggregation (34, 42).

In contrast to the redundant function of *ibpA* and *ibpB* in K-12 strains, we see in this work that disruption of either *ibpA* or *ibpB* causes severe growth defects at 45°C in the *E. coli* C122 strain (Fig. 1B-D). Although not an identical stress to heat shock, we have previously shown that *dnaK* and *clpB* genes are upregulated in C122 undergoing ϕX174 infection, with *dnaK* upregulated 2.5-fold and *clpB* upregulated 2.2-fold (10). Our inability to completely disrupt both *ibpA* and *ibpB* simultaneously in C122 without compensatory mutations (Fig. 2) may suggest that the *ibpAB* genes display synthetic sickness or lethality characteristics in this strain that are not present in K-12 strains. Other work by Rousset et al. (7) fits with this hypothesis, as they have shown that there are significant differences in gene essentiality between *E. coli* strains.

Another possibility is that cryptic recombination may have occurred in some previous reports and this was not detected because confirmation of mutants was done by PCR product sizing or similar low-resolution methods. With the advent of whole-genome sequencing, previously invisible confounding events such as the sequence duplication we observed (Fig. 2) may partially explain the differences we see to past work.

Despite the sHsps being highly upregulated during many different phage infections (10, 12, 56, 57) we have shown in this work that they do not appear to play a role in the *Microviridae* phage ϕX174 replication cycle. Previously, Young, et al (1989) found that lysis sensitivity increased in *E. coli* when plasmid encoded ϕX174 E protein and heat-shock genes *dnaK, dnaJ, groEL, and grpE* were present (24). Additionally, other folding chaperones were shown to be important for successful phage replication in phage PRD1, HK97, and T4, which all rely on the GroES/EL chaperones (20, 21, 67). In other phage infections, the *IbpA/B* genes were upregulated to some of the highest levels of any transcript or protein in *P. aeruginosa* and *E. coli* during PRR1 phage infection (57), along with other σ^32^ regulated genes (56), but they did not investigate the essentiality of these two sHSPs for phage infection.

Although more typically associated with cellular stress due to protein overexpression and increased temperatures (25), the sHsps appear to have another major role in maintaining membrane stability, which has been demonstrated in both prokaryotes and eukaryotes. In cyanobacterium *Synechocystis* sp. PCC 6803, HspA (or Hsp17) was found specifically associated with the thylakoid membrane under heat stress and functioned to reduce fluidity back to more normal levels (68–70). Under salt stress, increased expression of GroEL/GroES was observed in a *Synechocystis* mutant that lacked *hspA*; GroEL/GroES are also known to assist the folding of membrane-associated proteins and stabilize the lipid membrane structure (71, 72). Similarly, the Lo18 sHsp from the lactic acid bacterium *Oenococcus oeni* was found associated with the cell membrane because of increased membrane disorder, rather than cellular protein aggregation (73). *In* vitro studies have also shown that Lo18 interacts with liposomes and increases the molecular order of the lipid bilayer (74, 75). In *Lactobacillus plantarum*, deletion of *hsp 18*.*55* caused increased membrane fluidity when exposed to ethanol shock, whereas *L. plantarum* overexpressing Hsp18.55 had significant reduction in membrane fluidity compared to the wild-type strain (76). Functional characterisation of Hsp20 from the thermoacidophilic crenarchaeon *Sulfolobus acidocaldarius* has revealed that sHsps stabilise archaeal and bacterial membrane possibly through hydrophobic interactions with membrane lipids (archaeosomes and liposomes) (77). Furthermore, HSP17 (a sHsp from *Caenorhabditis elegans*), when heterologously expressed in *E. coli*, was found to be partially localised to periplasmic space and associated with inner membrane to maintain cell envelope integrity when *E. coli* were grown at lethal temperature (50°C) (78). The IbpA and IbpB proteins were previously found associated with the outer membrane in *E. coli* (30, 33) and are known to form a complex with lipoprotein NlpI and localise to the outer membrane during cell division (59).

One possible explanation for our results that show *ibpAB* is dispensable for ϕX174 replication, is that IbpA/B may be upregulated in response to both the large increase in protein production due to phage infection as well as changes to the membrane integrity during infection. A potentially similar response is seen with the Phage Shock Protein (Psp) complex which repairs cell envelope integrity, and is upregulated in both ϕX174 and PRD1 infections of *E. coli* (10, 56, 79, 80). Phage ϕX174 expresses a single lysis protein E during infection and its mechanism of action is thought to be through inhibiting a key enzyme (MraY) in cell wall peptidoglycan synthesis (81). We speculate that the progressive disruption to peptidoglycan structure and cell envelope integrity results in a signal that causes the upregulation of the *ibpAB* genes. The produced proteins may provide some transient protection to cell membrane integrity prior to lysis, but they are ultimately overwhelmed by lysis protein production and burst of phage progeny.

## Materials and Methods

### Bacterial strains, phage, plasmids, and culture conditions

Bacteria were routinely grown in lysogeny broth (LB) (Miller) medium (1% w/v tryptone, 0.5% w/v yeast extract, and 1% NaCl) and incubated at 37°C in an Infors MT multitron pro shaker at 240 RPM rotating orbitally at a 25 mm diameter. LB (Miller) agar (1.5% w/v) was used as solid medium. Unless otherwise stated, carbenicillin was used at 50 µg/mL, chloramphenicol was used at 20 µg/mL, and spectinomycin was used at 50 µg/mL. For routine ϕX174 phage amplification, a single plaque was added to mid-exponential phase culture of bacteria in phage-LB (LB supplemented with 2 mM CaCl_2_), and phage were purified as previously described (82). Phage titers were measured by the double-layer overlay method (83). Growth media, buffers, and reagents were sterilised by autoclaving and/or filtering through 0.2 µm membrane prior to use in the experiments.

The bacterial strains, phage, and plasmids used in this study are presented in Table 1. Synthetic DNA was acquired from Integrated DNA Technologies (Coralville, IA, USA), primers from Azenta Life Sciences (South Plainfield, NJ, USA), and they are described here (<u>Supplemental Table S1</u>).

### CRISPR/Cas9 mediated gene knockouts

Custom sgRNA plasmids, knockouts, and plasmid curing were performed as described (84, 85), except for transformation. Transformations were performed with chemically competent cells using the *Mix and Go! E. coli* Transformation and Buffer Set (Zymo Research, USA). Primers, sgRNA sequences, and donor DNA are described here (<u>Supplemental Table S1</u>). Confirmation of incorporated targeting sgRNA sequences and knockouts were performed through PCR amplification of targeted regions, followed by PCR product purification using the GenElute™ PCR Clean-Up Kit (Sigma Aldrich, U.S.A). All sequencing throughout was performed using the Macrogen sequencing service (Macrogen, South Korea).

Whole genome sequencing of C122, C122Δ*ibpA*, C122Δ*ibpB*, and C122Δ*ibpAB* strains was performed by preparing by Illumina sequencing libraries from gDNA samples using Nextera DNA Flex Library Preparation Kit. Libraries were sequenced on an Illumina MiSeq instrument (Sequencing @ UTS). Read processing and mapping was performed using Geneious Prime software.

### Assembly of plasmid for trans expression of ibpAB genes

The native *ibpAB* operon (NZ_LT906474 coordinates: 16,177-17,263) was assembly into linearized pULTRA-CNF plasmid using NEBuilder Assembly (New England Biolabs), resulting in pULTRA::*ibpAB*. A version of pULTRA-CNF lacking the inserted gene (pULTRA::empty) was also created (Table 1). Sequence-confirmed plasmids were transformation into C122Δ*IbpAB* and wild-type C122 using the *Mix and Go! E. coli* Transformation and Buffer Set (Zymo Research, U.S.A). Transformants were selected and grown in phage-LB under a selective marker spectinomycin (50 μg/mL).

### CRISPRi plasmid construction

The dCas9-sgRNA plasmid expression system used for CRISPRi, pFR56, harbors a constitutively expressed sgRNA and *dcas9* under the control of a DAPG-inducible PhlF promoter (7). We predicted all possible gRNAs targeting *ibpA* or *ibpB* using the CRISPR Guide RNA Design tool (https://crispr-browser.pasteur.cloud/guide-rna-design) (86). The gRNA sequence with the highest score on the non-template (coding) strand was selected, because binding of gRNA to non-template strand has been demonstrated more efficient in repressing transcription than binding to the template strand (6, 53). The selected gRNAs targeting *ibpA* or *ibpB* were cloned on pFR56 using a pair of divergent primers (87) (<u>Supplemental Table S1</u>), and the assembled gRNA plasmids were transformed into chemically competent NEB Turbo *E. coli* cells (New England Biolabs, C2984H). Transformants were selected on LB-Cm agar plates and PCR amplification of targeted regions was used to confirm correct plasmid assembly, followed by Sanger sequencing (Macrogen, South Korea). Sequence-verified plasmids were transformed into chemically competent C122, C122Δ*ibpA*, and C122Δ*ibpB* cells prepared by the Mix and Go! *E. coli* Transformation and Buffer Set (Zymo Research, USA).

### Bacterial growth assay

Isolates of *E. coli* C122 were grown overnight in 4 mL of LB at 37°C in an Infors MT multitron pro shaker at 250 rpm rotating orbitally at a 25 mm diameter. Overnight cultures were diluted 100-fold in 200 µL of LB or LB-Cm in a 96-well microtiter plate (Cellstar, 655180). The inoculates were grown at 30°C, 37°C, or 45°C/237 rpm (double orbital) for 20 h on a Biotek Synergy H1 microplate reader (Biotek, USA). The optical density at 600 nm (OD_600_) was measured every 5 min. The kinetic data were collected by Gen5 (version: 3.08) (Agilent, USA). The assay was performed in biological triplicates. The OD_600_ values as a function of time were plotted to generate the growth curve. The growth rate and the maximum population size at stationary phase were calculated using Growthcurver R package (88).

### In vitro bacterial killing assay

Bacterial growth and OD_600_ monitoring were described as above. When OD_600_ reached ~0.15, DAPG was added to the cultures to a final concentration of 50 µM to induce the expression of dCas9. Bacterial strains were allowed to grow for a further 40 minutes before infection with ϕX174 phage at MOI=5 and 0.001. In the control experiment, 1 µL carrier (70% (v/v) ethanol) was added as mock induction, and 2 µL phage-LB was added as mock phage infection. The assay was performed in biological triplicates. The virulence of ϕX174 against *E. coli* C122 strains was assessed by the liquid assay score as previously described, which is equal to the area between the control curve and the phage treatment curve; a score of zero indicate no inhibition of bacterial growth, whereas complete absence of bacterial growth of phage-infected culture results in a score of 100 (47). The area under each curve was calculated using Growthcurver R package (88).

### Efficiency of centre of infection (ECOI) assay

Centre of infection assays were conducted as previously described (89). Overnight bacterial cultures were diluted 100-fold in 10 mL of phage-LB or phage-LB-Cm, and were grown at 37°C/240 rpm. The OD_600_ was monitored in a spectrophotometer (UV-1280, Shimadzu, Japan). DAPG was added to a final concentration of 50 µM when OD_600_ reached ~0.3, and bacteria continued to grow for a further 40 min. Subsequently, 2 mL of bacterial culture was centrifuged and resuspended in 500 µL of phage-LB. To achieve an MOI of 0.001 or less, 60 pfu phage lysates were added. Phage were allowed to adsorb for 5 min at 37°C/900 rpm in a thermomixer (Eppendorf). The cells were placed on ice for 5 min and then washed twice with 500 µL of phage-LB to remove free phage. The cells with adsorbed phage were co-plated with 200 µL of wild type *E. coli* C122 overnight culture following the standard soft agar overlay method (83), and incubated overnight at 37°C. Resulting plaques were enumerated and the efficiency of centre of infection (ECOI) was calculated. The experiment was performed in four biological replicates.

### RNA isolation and RT-qPCR analysis

Biological triplicates of *E. coli* C122 were grown in 1.45 mL of phage-LB or phage-LB-Cm in a 96-well deep-well plate at 37°C in an Infors MT multitron pro shaker at 240 rpm rotating orbitally at a 25 mm diameter. Expression of dCas9 was induced at early exponential phase by DAPG to a final concentration of 50 µM. The bacteria were further grown for 40 min before infection with wild type ϕX174 at an MOI of 5. At 60 min after infection, samples were pelleted by centrifugation at 5,000 RCF for 10 min at 4°C and resuspended in 250 µL ice-cold 1x phosphate-buffered saline (PBS) at pH 7.4, RNAprotect bacterial reagent (Qiagen, 76506) was added and the cells treated according the manufacturer instructions. Following protection, RNA was isolated using the RNeasy Mini Kit (Qiagen, 74106) with the optional DNase step according to the manufacturer instructions.

The quality and quantity of RNA were determined using a NanoDrop spectrophotometer (NanoDrop One, Thermo Fisher Scientific, Waltham, MA, USA). All RNAs were reverse transcribed into cDNA using the High Capacity cDNA Reverse Transcription Kit (ThermoFisher Scientific, 4368814). RT-qPCR was performed with the LightCycler 480 SYBR Green I Master (Roche, 04707516001) in a LightCycler 480 II (Roche, USA) following manufacturer’s instructions in 10 µL volumes with an annealing temperature of 64°C. RT-qPCR was performed in technical triplicate and biological triplicate. Relative gene expression was computed using the comparative C_T_ method (90) after normalization to the *cysG* housekeeping gene with the infected samples as the treated samples and the uninfected the untreated.

### Statistical analysis

All statistical analyses were conducted with R program language (version 4.2.1) (https://cran.r-project.org/bin/macosx/big-sur-arm64/base/R-4.2.1-arm64.pkg). Student’s two-tailed t-test were performed to assess the differences in bacterial growth rate, liquid assay scores, ECOI of ϕX174, and fold changes of *ibpA/B* expression levels.

## Acknowledgements

We recognize that this research and analysis was conducted on the traditional lands of the Wallumattagal clan of the Dharug nation, the Gadigal, Wangal, and Cameraygal peoples of the Eora Nation. We thank Dr. François Rousset and David Bikard lab for providing the pFR56 plasmid.

B.W.W was supported by Macquarie Research Excellence Scholarship. H.X.Z was supported by the Australian Government’s Research Training Program (RTP) Scholarship. D.Y.L. was supported by the Macquarie University COVID Recovery Postdoctoral Fellowship. P.R.J was supported by NHMRC Ideas Grant APP1185399.

The contributions of authors of this work according to the CRediT contribution taxonomy were as follows. H.X.Z.: Formal analysis, Investigation, Methodology, Writing – original draft, Writing – review & editing; B.W.W.: Conceptualization, Formal analysis, Investigation, Methodology, Writing – original draft, Writing – review & editing; D.Y.L.: Formal analysis, Investigation, Methodology, Visualization, Writing – original draft, Writing – review & editing; M.P.M.: Funding acquisition, Project administration, Supervision, Writing – review & editing; P.R.J.: Conceptualization, Funding acquisition, Project administration, Resources, Supervision, Visualization, Writing - review & editing.

